# UTMOST, A Novel Single And Cross-Tissue TWAS (Transcriptome Wide Association Study), Reveals New ASD (Autism Spectrum Disorder) Associated Genes

**DOI:** 10.1101/2020.06.11.145524

**Authors:** Cristina Rodriguez-Fontenla, Angel Carracedo

## Abstract

**Background:** ASD is a complex neurodevelopmental disorder which may significantly impact on the affected individual’s life. Common variation (SNPs) could explain about 50% of ASD heritability. Despite this fact and the large size of the last GWAS meta-analysis, it is believed that hundreds of risk genes in ASD have yet to be discovered. New tools such as TWAS which integrate tissue-expression and genetic data, are a great approach to identify new ASD susceptibility genes.

**Methods:** The main goal of this study is to use UTMOST with the public available summary statistics from the largest ASD GWAS meta-analysis as genetic input. In addition, an *in silico* biological characterization for the novel associated loci was performed.

**Results:** Our results have demonstrated the association of 4 genes at the brain level (*CIPC, PINX1, NKX2-2* and *PTPRE*) and highlight the association of *NKX2-2, MANBA, ERI1* and *MITF* at the gastrointestinal level. This last finding is quite relevant given the well-established but unexplored relationship between ASD and gastrointestinal symptoms. Cross-tissue analysis has shown the association of *NKX2-2* and *BLK*.

**Conclusions:** UTMOST associated genes together with their *in silico* biological characterization seems to point to different biological mechanisms underlying ASD etiology not only restricted to brain tissue and involving other body tissues such as the gastrointestinal tissue.

## INTRODUCTION

ASD (Autism Spectrum Disorders) includes a range of NDDs (Neurodevelopmental Disorders) with onset in early development that are characterized by deficits in communication and social interactions, as well as by repetitive patterns of behavior and restrictive interests ^1^. ASD is a complex genetic disorder, involving both environmental and genetic factors. Although an important part of the genetic architecture of ASD is unknown, it is considered that thousands of genes may be involved even most of them remain unidentified and functionally uncharacterized. Rare genetic variation only explains 3% of ASD genetic risk even if it confers a high individual risk ^2^. However, common variation (SNPs; *single nucleotide polymorphisms*) could explain about 50% of ASD heritability. The most recent and the largest ASD GWAS meta-analysis done until now, including 18,381 ASD cases and 27,969 controls, has reported 93 genome-wide significant markers in 3 separate loci (top SNP: rs910805; p-value: 2.04 × 10^−9^)^3^. Another methodological approach for common variation are gene-based association analysis (GBA) methods that employ the p-values for each SNP within a gene to obtain a single statistic at this level. Thus, MAGMA has identified 15 genes, most of them located near the genome-wide significant SNPs identified in the GWAS, but 7 genes reveal associatio in 4 additional loci (*KCNN2, MMP12, NTM* and a cluster of genes on chromosome 17)^3^. Additional GBA methods using other algorithms as PASCAL have helped to define the association of other genes located in the same LD region than those found by MAGMA (*NKX2-4, NKX2-2, CRHR1-IT1, C8orf74* and *LOC644172*)^4^ In addition to GBA, bioinformatic approaches that integrate functional data are increasingly used to highlight new genes underlying GWAS summary statistics. Transcriptome-wide association studies (TWAS) have emerged as useful tools to study the genetic architecture of complex traits. Among them, MetaXcan^5^ and FUSION^6^ are well-known TWAS methods.

UTMOST (Unified Test for MOlecular SignaTures) has been recently reported as a novel framework for single and cross-tissue expression imputation. UTMOST is able to consider the joint effect of SNPs (summary statistics) across LD regions (1000 Genomes) and to integrate tissue expression data (GTeX) creating single and cross-tissue covariance matrices that will help to define the gene-trait associations. UTMOST performance was demonstrated at several levels and its accuracy was also well proved as it was able to identify a greater number of associations in biologically relevant tissues for complex diseases ^7^.

The main aim of this paper is to further mine the summary data from the ASD meta-analysis using UTMOST. In addition, an *in silico* biological characterization for the novel associated loci will be carried out using bioinformatic approaches (DEG, pathway, gene network and an exploratory enhancer analysis).

Overall, our results have demonstrated the association of *CIPC, PINX1, NKX2-2* and *PTPRE* at the brain level and have also revealed the relevance of gastrointestinal tissue in ASD etiology through the association of other genes (*NKX2-2, MANBA, ERI1* and *MITF)*.

## METHODS AND MATERIALS

### 1. Datasets

Summary statistics from the latest ASD GWAS meta-analysis were obtained from the public repository available in the PGC website (http://www.med.unc.edu/pgc/results-and-downloads). The following data set was employed: iPSYCH_PGC_ASD_Nov2017.gz (Grove et al., 2019) which includes the meta-analysis of ASD by the Lundbeck Foundation Initiative for Integrative Psychiatric Research (iPSYCH) and the Psychiatric Genomics Consortium (PGC) released in November 2017. The data set comprises a total of 18,381 cases and 27,969 controls. Additional information about the genotyping and QC methods employed are available at the PGC website.

### 2. TWAS Analysis using UTMOST

UTMOST^7^ (https://github.com/Joker-Jerome/UTMOST) was run as a single tissue association test for 44 GTeX tissues (single_tissue_association_test.py) and a cross tissue association test combining gene-trait associations was run by the joint GBJ test (joint_GBJ_test.py). Both tests use the previous summary statistics of the ASD GWAS meta-analysis as an input^3^. UTMOST pre-calculated covariance matrices for single-tissue (covariance_tissue/) and joint test (covariance_joint/) were downloaded. Other neccesary command parameters were used by default. Transcriptome-wide significance for single tissue analysis was stablished as p-value =3.85 × 10 ^-6^ (0.05/12984(maximum number of genes tested) for brain tissues and p-value=3.42 × 10 ^-6^ (0.05/14586 (maximum number of genes tested) for non-brain tissues after Bonferroni correction. Transcriptome-wide significance for joint test was established as p-value= 3.27 × 10 ^-6^ (0.05/15274) considering the number of effective test (Table 1, 2 & 3) (Supplementary .csv files for each tissue). Covariances matrices are only available for the 44 tissues of GteX v6: Adipose Subcutaneous, Adipose Visceral Omentum, Adrenal Gland, Artery Aorta, Artery Coronary, Artery Tibial, Brain Anterior cingulate cortex BA24, Brain Caudate basal ganglia, Brain Cerebellar Hemisphere, Brain Cerebellum, Brain Cortex, Brain Frontal Cortex BA9, Brain Hippocampus, Brain Hypothalamus, Brain Nucleus accumbens basal ganglia, Brain Putamen Basal ganglia, Breast Mammary Tissue, Cells EBV-transformed lymphocytes, Cells Transformed fibroblasts, Colon Sigmoid, Colon Transverse, Esophagus Gastroesophageal Junction, Esophagus Mucosa, Esophagus Muscularis, Heart Atrial Appendage, Heart Left Ventricle Liver, Lung Muscle Skeletal, Nerve Tibial, Ovary, Pancreas, Pituitary, Prostate, Skin Not Sun Exposed Suprapubic, Skin Sun Exposed Lower Leg, Small Intestine, Terminal Ileum, Spleen, Stomach, Testis, Thyroid, Uterus, Vagina, Whole Blood. Morpheus software (https://software.broadinstitute.org/morpheus/) was used to display the Z scores of UTMOST significant genes across GTeX Brain tissues. UTMOST significance as a *Z* score in brain tissues is **∼**4.6. Grey squares in the heatmap indicate that the gene weights were not available in the target tissue (Figure 1).

**Table 1.**
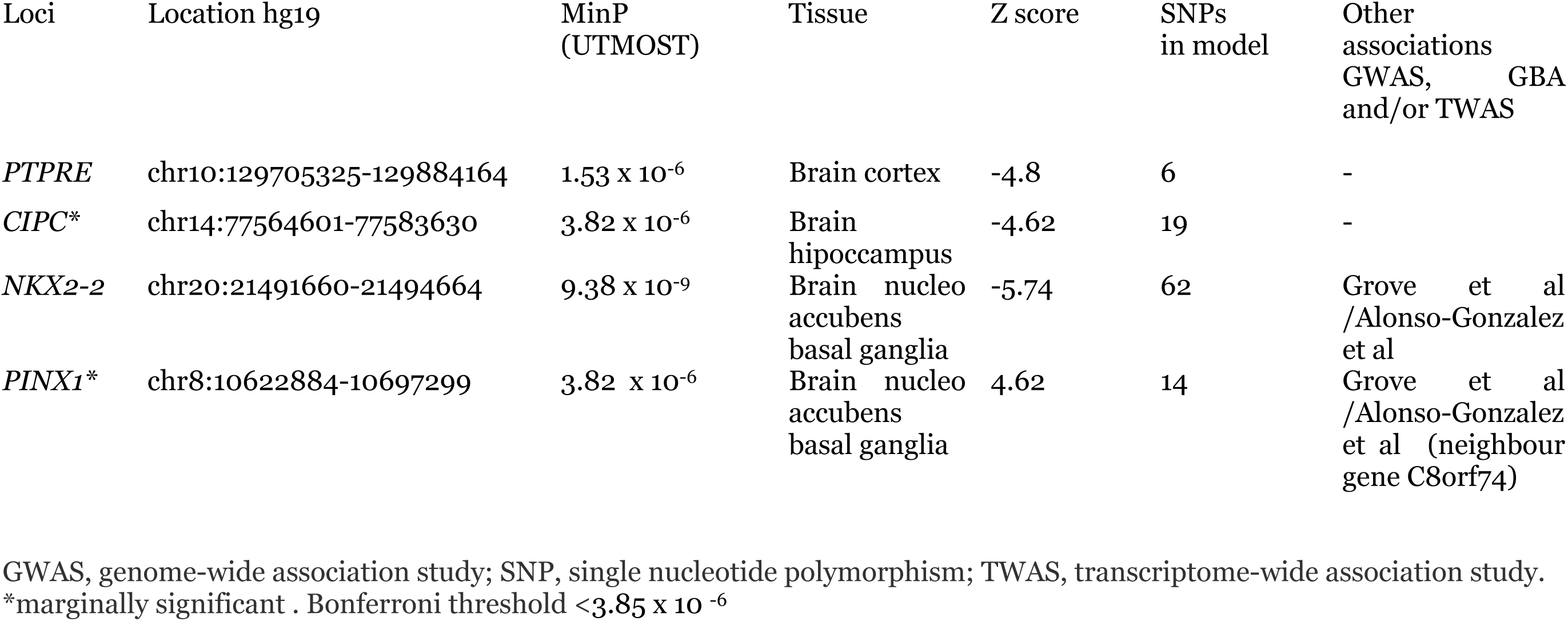
UTMOST Single tissue analysis (Brain tissues). List of Independent Significant Loci.

| Loci | Location hg19 | MinP<br>(UTMOST) | Tissue | Z score | SNPs<br>in model | Other<br>associations<br>GWAS, GBA<br>and/or TWAS |
| --- | --- | --- | --- | --- | --- | --- |
| *PTPRE* | chr10:129705325-129884164 | 1.53 x 10<sup>-6</sup> | Brain cortex | -4.8 | 6 | - |
| *CIPC\** | chr14:77564601-77583630 | 3.82 x 10<sup>-6</sup> | Brain<br>hippocampus | -4.62 | 19 | - |
| *NKX2-2* | chr20:21491660-21494664 | 9.38 x 10<sup>-9</sup> | Brain nucleo<br>accubens<br>basal ganglia | -5.74 | 62 | Grove et al<br>/Alonso-Gonzalez<br>et al |
| *PINX1\** | chr8:10622884-10697299 | 3.82 x 10<sup>-6</sup> | Brain nucleo<br>accubens<br>basal ganglia | 4.62 | 14 | Grove et al<br>/Alonso-Gonzalez<br>et al (neighbour<br>gene C8orf74) |

GWAS, genome-wide association study; SNP, single nucleotide polymorphism; TWAS, transcriptome-wide association study.
\*marginally significant . Bonferroni threshold <3.85 x 10<sup>-6</sup>

**Table 2.** UTMOST Single tissue analysis (Non Brain tisues). List of Independent Significant Loci.

| Loci | Location hg19 | MinP<br>(UTMOST) | Tissue | Z<br>score | SNPs<br>in<br>model | Other<br>GWAS,<br>GBA and/or<br>TWAS |
| --- | --- | --- | --- | --- | --- | --- |
| ***NKX2-2*** * | chr20:21491660-21494664 | 5.28 x 10<sup>-6</sup> | Colon transverse | -4.55 | 2 | Grove et al<br>/Alonso-Gonzalez et al |
| ***MANBA*** | chr4:103552643-103682151 | 3.2 x 10<sup>-6</sup> | Esophagus<br>muscularis | 4.66 | 47 | - |
| ***ERI1*** * | chr8:8860314-8890849 | 3.84 x 10<sup>-6</sup> | Esophagus<br>muscularis | 4.62 | 124 | - |
| *DOK5* | chr20:53092266-53267710 | 4.86 x 10<sup>-7</sup> | Liver | -409.74 | 4 | - |
| *ATXN1* | chr6:16299343-16761721 | 2.4 x 10<sup>-6</sup> | Nerve tibial | -4.72 | 27 | -- |
| *FERMT2* | chr14:53323989-53417815 | 3.1 x 10<sup>-6</sup> | Skin not sun<br>exposed | -5.49 | 14 |  |
| ***MITF*** | chr3:69788586-70017488 | 2.34 x 10<sup>-6</sup> | Stomach | 4.72 | 111 | - |
| *CTSB* | chr8:11700034-11725646 | 3.36 x 10<sup>-6</sup> | Thyroid | 4.65 | 66 | - |
| *ANGEL1* | chr14:77253586-77279283 | 1.89 x 10<sup>-6</sup> | Uterus | 4.76 | 83 | - |

GWAS, genome-wide association study; SNP, single nucleotide polymorphism; TWAS, transcriptome-wide association study. marginally significant . \*Genes in bold are associated within gastrointestinal tissues. Bonferroni threshold = $3.42 \times 10^{-6}$

**Figure 1.**
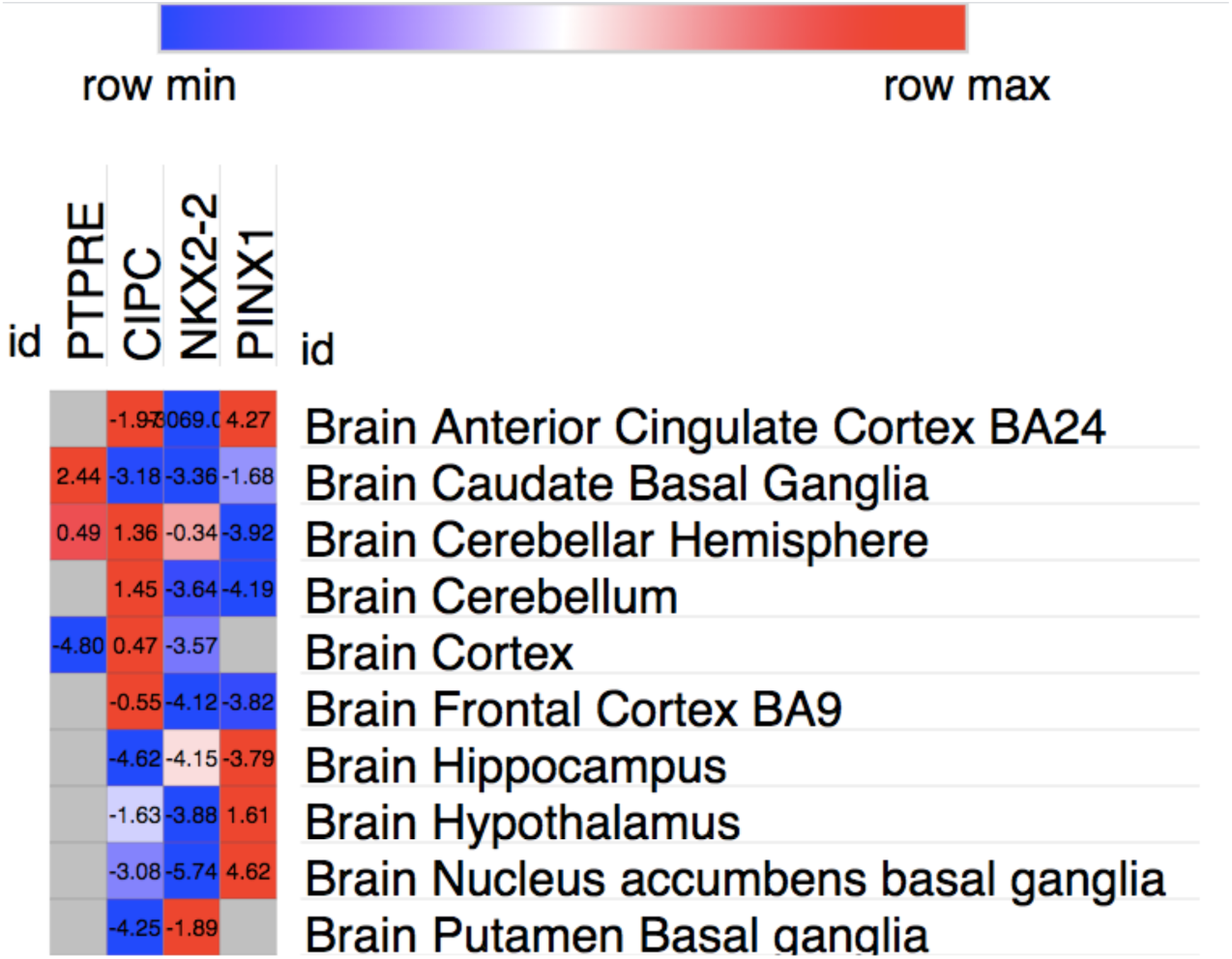
Heatmap representing UTMOST significant genes across weight sets and brain areas. UTMOST significance as a *Z* score is **∼**4.6. Grey squares indicate that gene weights were not available in the target tissue.

### 3. DEG analysis with BrainSpan

GENE2FUNC, a tool of FUMA^8^ (https://fuma.ctglab.nl/), was employed to carry out a gene expression heatmap and an enrichment analysis of differentially brain expressed genes (DEG) using BrainSpan RNA-seq data. Those genes represented in Table 4 were used as an input. Expression values are TPM (Transcripts Per Million) for GTEx v6 and RPKM (Read Per Kilobase per Million). Heatmaps display the normalized expression value (zero mean normalization of log2 transformed expression), and darker red means higher relative expression of that gene in each label, compared to a darker blue color in the same label. DEG analysis was carried out creating differentially expressed genes for each expression data set. In order to define DEG sets, two-sided Student’s t-test were performed per gene and per tissue against the remaining labels (tissue types or developmental stages). Those genes with a p-value < 0.05 after Bonferroni correction and a log fold change ≥ 0.58 are defined as DEG. The direction of expression was considered. The -log10 (p-value) refers to the probability of the hypergeometric test (Figure 2 and 3).

**Figure 2.**
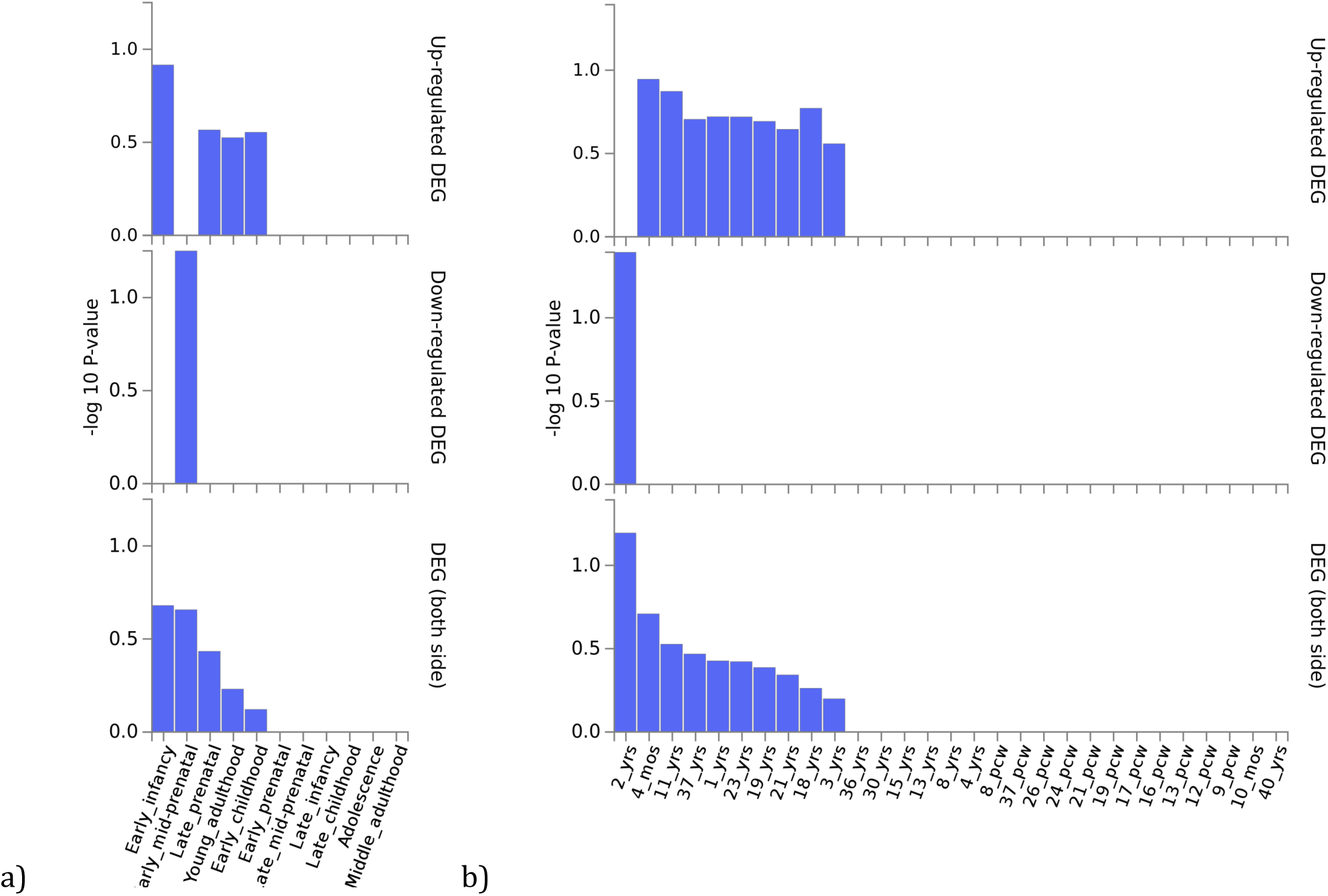
DEG analysis for *PTPRE, CIPC, PINX1* and *NKX2-2* using BrainSpan 11 general developmental stages of brain samples (a) and BrainSpan 29 different ages of brain data (b) There are not significantly enriched DEG sets.

**Figure 3.**
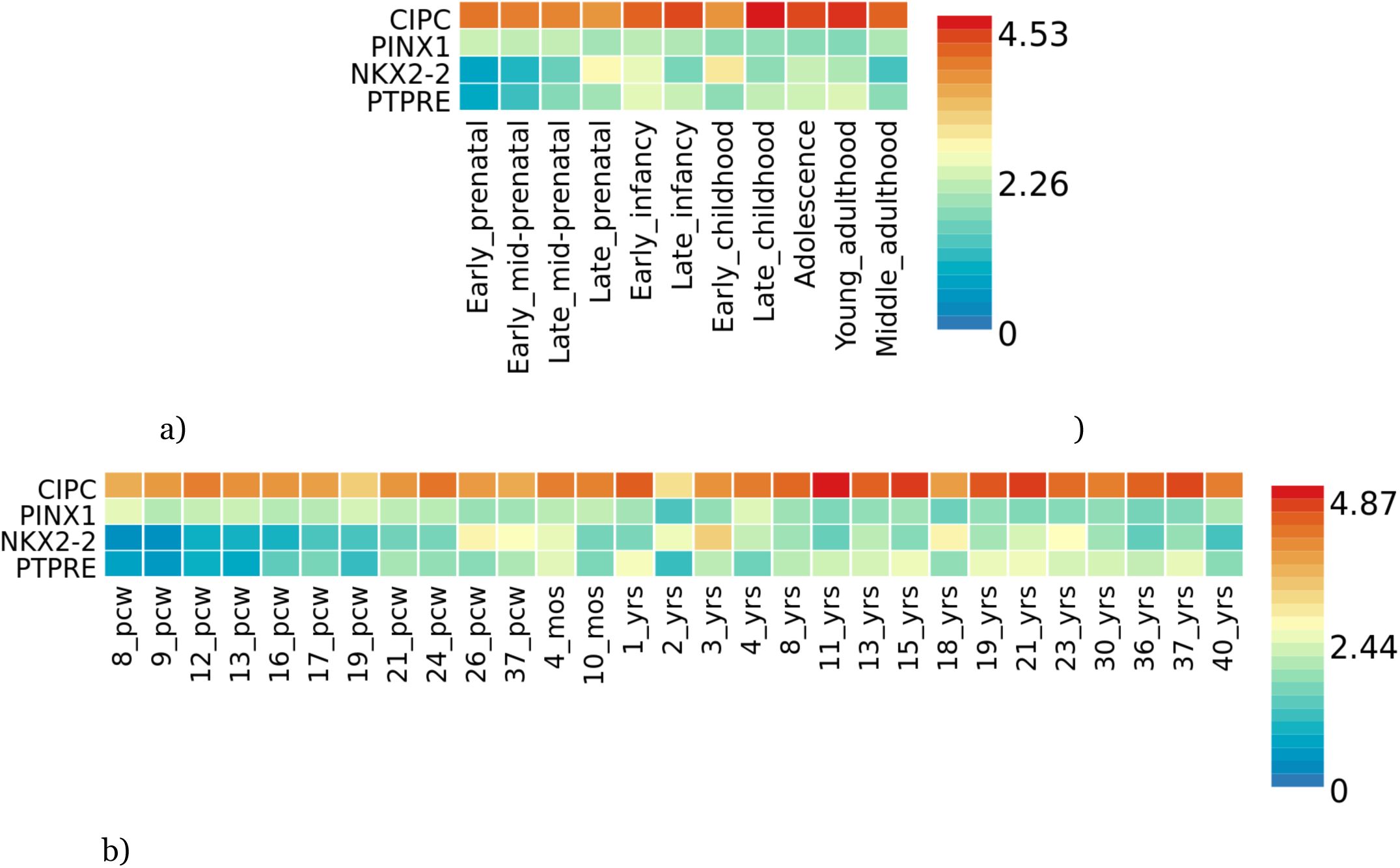
Gene expression heatmaps for *PTPRE, CIPC, PINX1* and *NKX2-2* using BrainSpan 11 general developmental stages of brain samples (a) and BrainSpan 29 different ages of brain data (b). Genes are ordered by expression clusters and brain samples and ages by alphabetical order.

### 4. GeneMANIA and Metascape analysis

GeneMANIA^9^(https://genemania.org/) was used to build a gene network for the UTMOST associated genes by the single tissue analysis (brain and gastrointestinal tissues) and by the joint tissue analysis (Table 4). Each gene-network was subsequently analyzed with Metascape (https://metascape.org/) ^10^ to carry out a pathway enrichment and a protein-protein interaction enrichment using the Evidence Counting (GPEC) prioritization tool. For each given gene list, pathway and process enrichment analysis has been carried out with the following ontology sources: GO Biological Processes, GO Cellular Components and GO Molecular Functions. The enrichment background includes all the genes in the genome. Terms with a p-value < 0.01, a minimum count of 3, and an enrichment factor > 1.5 (the enrichment factor is the ratio between the observed counts and the counts expected by chance) are collected and grouped into clusters based on their membership similarities. More specifically, p-values are calculated based on the accumulative hypergeometric distribution, and q-values are calculated using the Benjamini-Hochberg procedure to account for multiple testings. Kappa scores are used as the similarity metric when performing hierachical clustering on the enriched terms, and sub-trees with a similarity of > 0.3 are considered a cluster. We select the terms with the best p-values from each of the 20 clusters, with the constraint that there are no more than 15 terms per cluster and no more than 250 terms in total (Figures 4, 5, 6). The network is visualized using Cytoscape, where each node represents an enriched term and is colored first by its cluster ID (or each given gene list, protein-protein interaction enrichment analysis has been carried out with the following databases: BioGrid6, InWeb_IM7, OmniPath8. The resultant network contains the subset of proteins that form physical interactions with at least one other member in the list. If the network contains between 3 and 500 proteins, the Molecular Complex Detection (MCODE) algorithm has been applied to identify densely connected network components. The MCODE networks identified for individual gene lists are shown in Figures 4, 5, 6.

**Figure 4.**
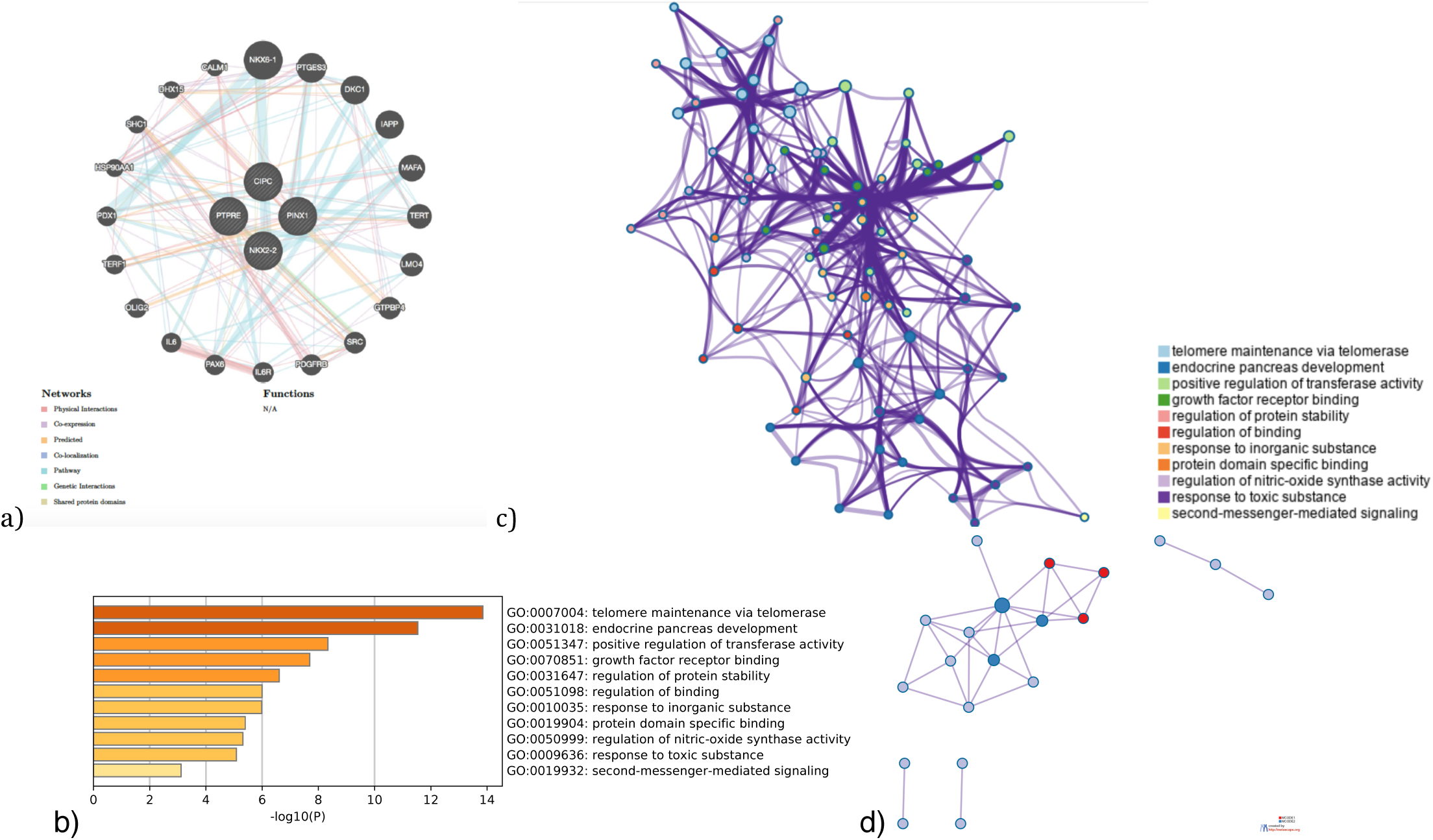
Gene networks, Enriched Ontology Clusters and PPI interaction analysis fot the Brain associated genes (*CIPC, PINX1, NKX2-2* and *PTPRE*) by UTMOST (Single tissue analysis). a) GeneMania gene network, b) Bar graph of enriched terms across input gene lists, colored by p-values, c) Network of enriched terms: colored by cluster ID, where nodes that share the same cluster ID are typically close to each other, d) Protein-protein interaction network and MCODE components identified in the gene lists. (MCODE1:red)

**Figure 5.**
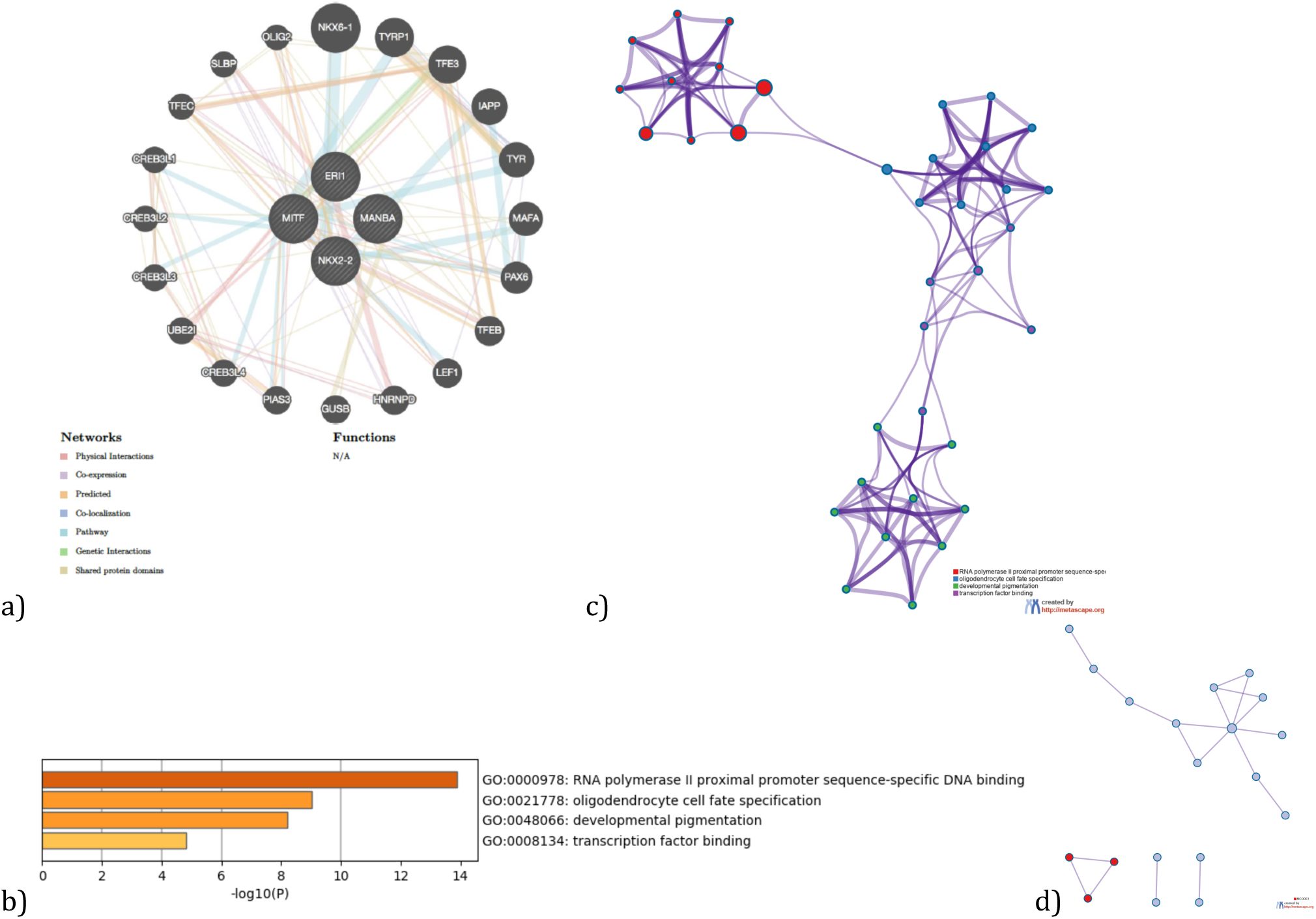
Gene networks, Enriched Ontology Clusters for the associated genes by UTMOST Single Tissue Analysis (gastrointestinal tissues) and their predicted interactors (*NKX2-2, MANBA, ERI1, MITF*). a) GeneMania gene network, b) Bar graph of enriched terms across input gene lists, colored by p-values, c) Network of enriched terms: colored by cluster ID, where nodes that share the same cluster ID are typically close to each other.

**Figure 6.**
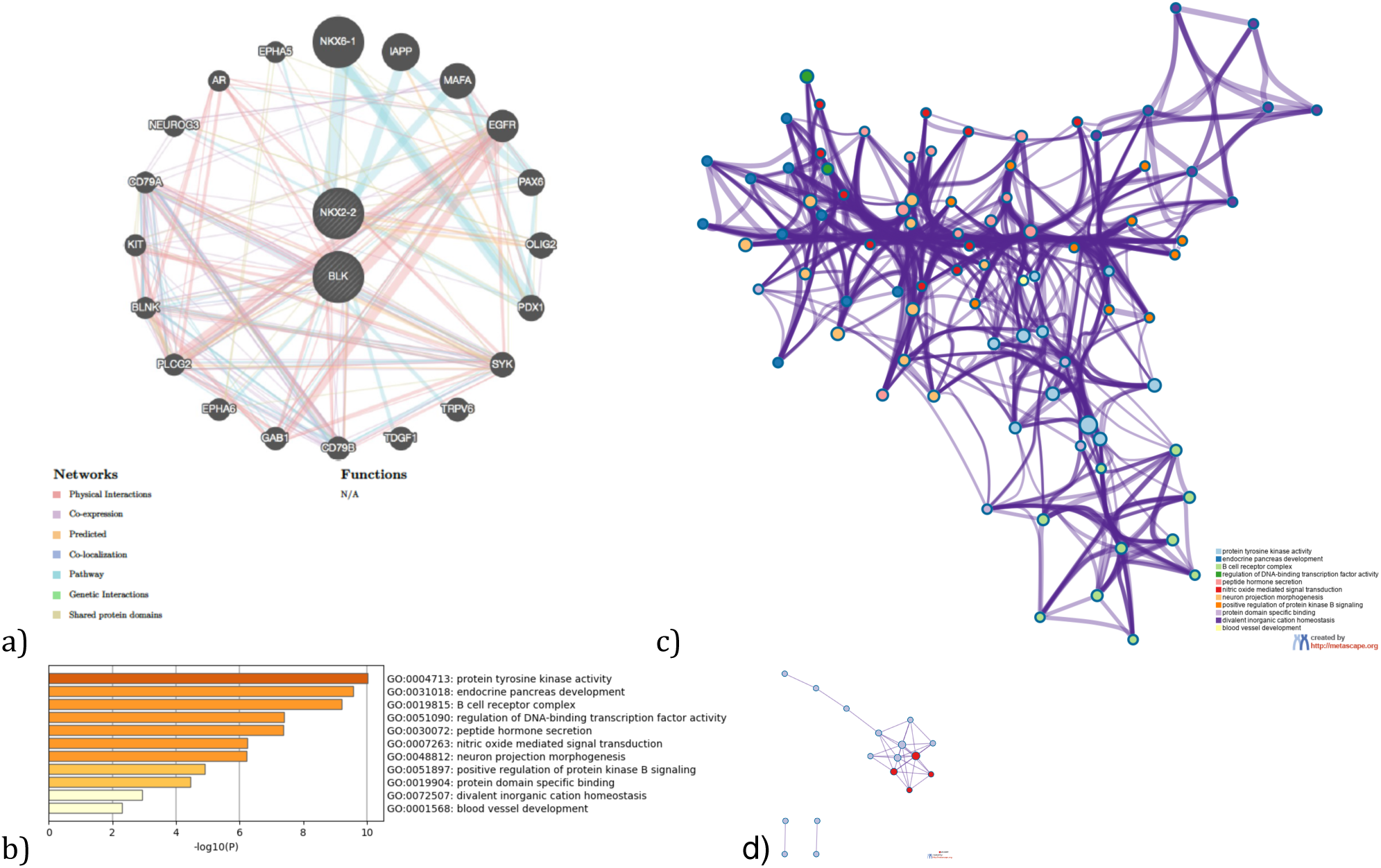
Gene network, Enriched Ontology clusters and PPI interaction analysis for the associated genes by UTMOST Cross Tissue Analysis (*BLK* and *NKX2-2*) and their interactors.). a) GeneMania gene network b) Bar graph of enriched terms across input gene lists, colored by p-values. c) Network of enriched terms: colored by cluster ID, where nodes that share the same cluster ID are typically close to each other. d) Protein-protein interaction network and MCODE components identified in the gene lists (MCODE1:red).

### 5. Enhancer analysis (dbSUPER)

dbSUPER (https://asntech.org/dbsuper/) was used to perform an exploratory enhancer analysis for the UTMOST associated genes (single-tissue analysis) and for those tissues (brain and gastrointestinal) available at dbSUPER. The parameters selected were: SEs ranking method (H3K27ac), the peak calling was done with MACS (version 1.4.1) with parameters -p 1e-9,-keep-dup = auto, -w - S -space = 50, the sitching distance was established at 12,5 kb, the TSS exclusive zone was set at +/-2 kb and the enhancer gene assignment was done within a 50 kb window.

## RESULTS

### 1. UTMOST analyses and comparison with previous results

UTMOST Single Tissue Analysis (Brain tissues) showed association of two loci, *NKX2-2* and *PTPRE*, while other two loci, *CIPC* and *PINX1*, showed a marginal association according to the threshold established by Bonferroni (Table 1, Supplementary .csv files). *NKX2-2* was previously identified as an associated gene by two different GBA algorithms, MAGMA and PASCAL^3,4^. The result obtained with UTMOST also serves to indicate the nucleus accubens basal ganglia as the brain area in which it may be acting. Although the association of *PTPRE* was not obtained as such in previous analyses, the association of its neighboring gene *C8orf74* was noted in a previous study and one of the index SNPs in ASD’s latest GWAS is located near *PINX1* (rs10099100) (Supplementary Figure 1a, 1b) ^3,4^.

*NKX2-2* was also marginally associated by the UTMOST analysis of non-brain tissue together with other genes. It should be noted that some genes are tissue-specific for gastric and intestinal tissues such as stomach, esophagus and colon (*MANBA, ERI1, MITF, NKX2-2*). The association of *NKX2-2* in colon is noteworthy because this is a previous reported ASD risk gene that now is highlighted again by UTMOST in brain tissues. It seems that *NKX2-2* together with *BLK* may play a role in ASD etiology but not only at the brain level since UTMOST cross-tissue analysis also found association for both genes (Table2, Table 3, Supplementary Figure 1a, 2a, 2b, Supplementary .csv files) To evaluate the importance of each brain tissue in ASD etiology, a secondary analysis was performed using Z scores values for each ASD associated gene across GTeX brain tissues. The heatmap showed that there is a wide specificity of association in terms of Z score for each gene and brain tissue indicating the importance of conducting tissue specific analyses such as UTMOST (Figure 1).

**Table 3.** UTMOST Cross tissue analysis. List of Independent Significant Loci.

| Loci | Location hg19 | MinP<br>(UTMOST) | test<br>score | Other<br>associations<br>GWAS,<br>GBA<br>and/or<br>TWAS |
| --- | --- | --- | --- | --- |
| *BLK* | chr8:11351521-11422108 | 2.85 x 10<sup>-6</sup> | 12.3 | Grove et al |
| *NKX2-2* | chr20:21491660-21494664 | 3.7 x 10<sup>-7</sup> | 13.9 | Grove et al<br>/Alonso-Gonzalez et<br>al |

GWAS, genome-wide association study; SNP, single nucleotide polymorphism; TWAS, transcriptome-wide association study . Bonferroni threshold=3.27 x 10<sup>-6</sup>

### 2. DEG Analysis with GENE2FUNC tool (FUMA)

DEG Analysis with BrainSpan data (11 general developmental stages of brain samples and 29 different ages of brain samples) for the brain associated set of genes have not shown any significant result (Figure 2a, 2b). However, *CIPC* have shown overexpression across every single developmental stage in comparison with the remaining ASD associated genes (Figure 3a,3b).

### 3. GeneMANIA and Metascape Analysis

GeneMANIA was used to find out the possible interactors with the associated genes (Table 4). We have proposed three different analyses. One based on the associated genes in brain; another one based on the associated genes in gastrointestinal tissue, given the associations pointed out by UTMOST and the previous implications of gastrointestinal abnormalities and symptoms in ASD and the lack of biological knowledge about them and a final analysis focused on the genes identified by the cross-tissue analysis and their interactors. The general goal is to delineate the biological pathways underlying each group of genes and the differences between them.

**Table 4.** List of associated genes for each analysis (bold) and their predicted GeneMANIA interactors that were used as input for Metascape analysis.

| UTMOST Single Brain | UTMOST Single Gastrointestinal | UTMOST Cross analysis |
| --- | --- | --- |
| **CIPC**<br>**PINX1**<br>**NKX2-2**<br>**PTPRE**<br>NKX6-1<br>PTGES3<br>DKC1<br>IAPP<br>MAFA<br>TERT<br>LMO4<br>GTPBP4<br>SRC<br>PDGFRB<br>IL6R<br>PAX6<br>IL6<br>OLIG2<br>TERF1<br>PDX1<br>HSP90AA1<br>SHC1<br>DHX15<br>CALM1 | **NKX2-2**<br>**MANBA**<br>**ERI1**<br>**MITF**<br>NKX6-1<br>TYRP1<br>TFE3<br>IAPP<br>TYR<br>MAFA<br>PAX6<br>TFEB<br>LEF1<br>HNRNPD<br>GUSB<br>PIAS3<br>CREB3L4<br>UBE2I<br>CREB3L3<br>CREB3L2<br>CREB3L1<br>TFEC<br>SLBP<br>OLIG2 | **NKX2-2**<br>**BLK**<br>NKX6-1<br>IAPP<br>MAFA<br>EGFR<br>PAX6<br>OLIG2<br>PDX1<br>SYK<br>TRPV6<br>TDGF1<br>CD79B<br>GAB1<br>EPHA6<br>PLCG2<br>BLNK<br>KIT<br>CD79A<br>NEUROG3<br>AR<br>EPHA5 |

Gene network, enriched ontology clusters and PPI interaction analysis for the brain associated genes (*CIPC, PINX1, NKX2-2* and *PTPRE*) by UTMOST Single tissue analysis and their interactors highlights different biological pathways mainly involved in telomere maintenance and transcription regulation (Table 5, 6 and Figure 4). However, associated genes in gastrointestinal tissue (*NKX2-2, MANBA, ERI1* and *MITF*) and their interactors mainly regulate DNA transcription by RNA polymerase II and the fate of oligodendrocytes. This is an interesting finding given the possible involvement of these cells in the enteric nervous system in ASD (Table 7, 8 and Figure 5). Finally, *NKX2-2* and *BLK* both associated in the cross-tissue analysis, seem to point to very diverse biological pathways such as protein tyrosine kinase activity, B cell receptor complex, regulation of DNA-binding transcription factor activity and neuron projection morphogenesis, among others (Table 8,9 and Figure 6).

**Table 5.** Top 11 clusters with their representative enriched terms (one per cluster) for the associated genes (*CIPC PINX1, NKX2-2* and *PTPRE*) (Single tissue analysis (Brain)) and their interactors. “Count” is the number of genes in the user-provided lists with membership in the given ontology term. “%” is the percentage of all of the provided genes that are found in the given ontology term (only input genes with at least one ontology term annotation are included in the calculation). “Log10(P)” is the p-value in log base 10. “Log10(q)” is the multi-test adjusted p-value in log base 10.

| GO | Category | Description | Count | % | Log10(P) | Log10(q) |
| --- | --- | --- | --- | --- | --- | --- |
| GO:0007004 | GO Biological Processes | telomere maintenance via telomerase | 7 | 43.75 | -13.85 | -9.50 |
| GO:0031018 | GO Biological Processes | endocrine pancreas development | 6 | 26.09 | -11.54 | -8.33 |
| GO:0051347 | GO Biological Processes | positive regulation of transferase activity | 8 | 50.00 | -8.34 | -5.76 |
| GO:0070851 | GO Molecular Functions | growth factor receptor binding | 5 | 31.25 | -7.69 | -5.24 |
| GO:0031647 | GO Biological Processes | regulation of protein stability | 6 | 26.09 | -6.61 | -4.30 |
| GO:0051098 | GO Biological Processes | regulation of binding | 6 | 26.09 | -6.00 | -3.83 |
| GO:0010035 | GO Biological Processes | response to inorganic substance | 6 | 37.50 | -5.98 | -3.83 |
| GO:0019904 | GO Molecular Functions | protein domain specific binding | 6 | 37.50 | -5.40 | -3.36 |
| GO:0050999 | GO Biological Processes | regulation of nitric-oxide synthase activity | 3 | 18.75 | -5.31 | -3.29 |
| GO:0009636 | GO Biological Processes | response to toxic substance | 6 | 26.09 | -5.08 | -3.10 |
| GO:0019932 | GO Biological Processes | second-messenger-mediated signaling | 4 | 17.39 | -3.12 | -1.51 |

**Table 6.**
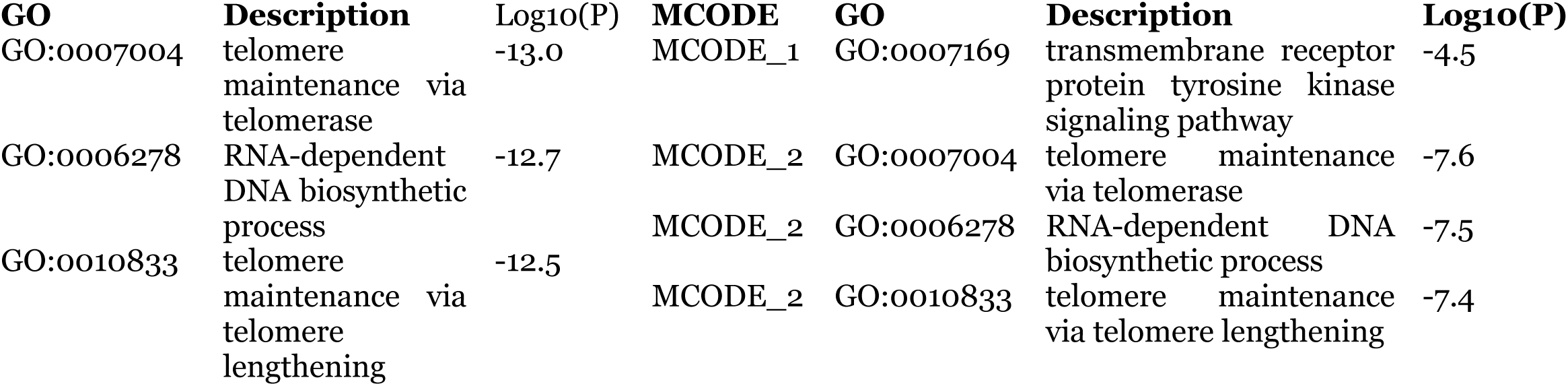
Protein-protein interaction network and MCODE components identified in the gene lists for the associated genes (*CIPC PINX1, NKX2-2* and *PTPRE*) (Single tissue analysis (Brain)) and their interactors

**Table 7.** Top 4 clusters with their representative enriched terms (one per cluster) fot the associated genes by UTMOST Single Tissue Analysis (GI tissues) (NKX2-2, MANBA, ERI1, MITF) and their predicted interactors. “Count” is the number of genes in the user-provided lists with membership in the given ontology term. “%” is the percentage of all of the provided genes that are found in the given ontology term (only input genes with at least one ontology term annotation are included in the calculation). “Log10(P)” is the p-value in log base 10. “Log10(q)” is the multi-test adjusted p-value in log base 10.

| **GO** | **Category** | **Description** | **Count** | **%** | **Log10(P)** | **Log10(q)** |
| --- | --- | --- | --- | --- | --- | --- |
| GO:0000978 | GO Molecular Functions | RNA polymerase II proximal promoter sequence-specific DNA binding | 12 | 57.14 | -13.89 | -9.63 |
| GO:0021778 | GO Biological Processes | oligodendrocyte cell fate specification | 3 | 25.00 | -9.03 | -5.82 |
| GO:0048066 | GO Biological Processes | developmental pigmentation | 4 | 33.33 | -8.21 | -5.12 |
| GO:0008134 | GO Molecular Functions | transcription factor binding | 6 | 28.57 | -4.81 | -2.47 |

**Table 8.** Protein-protein interaction network and MCODE components identified in the gene lists for the GI associated genes (and their interactors.

| GO | Description | Log10(P) | MCODE | GO | Description | Log10(P) |
| --- | --- | --- | --- | --- | --- | --- |
| GO:0000978 | RNA polymerase II proximal promoter sequence-specific DNA binding | -11.2 | MCODE_1 | GO:0035497 | cAMP response element binding | -9.8 |
| GO:0000987 | proximal promoter sequence-specific DNA binding | -11.0 | MCODE_1 | GO:0030968 | endoplasmic reticulum unfolded protein response | -6.9 |
| GO:0001228 | DNA-binding transcription activator activity, RNA polymerase II-specific | -10.8 | MCODE_1 | GO:0034620 | cellular response to unfolded protein | -6.7 |

**Table 9.** Top 11 clusters with their representative enriched terms (one per cluster) fot the associated genes by UTMOST Cross Tissue Analysis (*BLK* and *NKX2-2)* and their predicted interactors. “Count” is the number of genes in the user-provided lists with membership in the given ontology term. “%” is the percentage of all of the provided genes that are found in the given ontology term (only input genes with at least one ontology term annotation are included in the calculation). “Log10(P)” is the p-value in log base 10. “Log10(q)” is the multi-test adjusted p-value in log base 10.

| **GO** | **Category** | **Description** | **Count** | **%** | **Log10(P)** | **Log10(q)** |
| --- | --- | --- | --- | --- | --- | --- |
| GO:0004713 | GO Molecular Functions | protein tyrosine kinase activity | 6 | 42.86 | -10.03 | -5.68 |
| GO:0031018 | GO Biological Processes | endocrine pancreas development | 5 | 25.00 | -9.57 | -5.64 |
| GO:0019815 | GO Cellular Components | B cell receptor complex | 3 | 21.43 | -9.21 | -5.56 |
| GO:0051090 | GO Biological Processes | regulation of DNA-binding transcription factor activity | 7 | 35.00 | -7.40 | -4.63 |
| GO:0030072 | GO Biological Processes | peptide hormone secretion | 6 | 30.00 | -7.37 | -4.63 |
| GO:0007263 | GO Biological Processes | nitric oxide mediated signal transduction | 3 | 21.43 | -6.25 | -3.89 |
| GO:0048812 | GO Biological Processes | neuron projection morphogenesis | 7 | 35.00 | -6.21 | -3.87 |
| GO:0051897 | GO Biological Processes | positive regulation of protein kinase B signaling | 4 | 20.00 | -4.91 | -2.92 |
| GO:0019904 | GO Molecular Functions | protein domain specific binding | 5 | 35.71 | -4.45 | -2.58 |
| GO:0072507 | GO Biological Processes | divalent inorganic cation homeostasis | 4 | 18.18 | -2.92 | -1.26 |
| GO:0001568 | GO Biological Processes | blood vessel development | 4 | 18.18 | -2.31 | -0.69 |

**Table 10.** Protein-protein interaction network and MCODE components identified in the gene lists for the cross analysis associated genes and their interactors.

| GO | Description | Log10(P) | MCODE | GO | Description | Log10(P) |
| --- | --- | --- | --- | --- | --- | --- |
| GO:0030183 | B cell differentiation | -9.6 | MCODE_1 | GO:0030183 | B cell differentiation | -9.1 |
| GO:0004713 | protein tyrosine kinase activity | -9.4 |  |  |  |  |
| GO:0007169 | transmembrane receptor protein tyrosine kinase signaling pathway | -9.4 | MCODE_1 | GO:0042113 | B cell activation | -7.5 |
|  |  |  | MCODE_1 | GO:0030098 | lymphocyte differentiation | -7.3 |

### 4. Enhancer Analysis (dbSUPER)

Given the tissue specificity given by UTMOST associations, we found interesting to perform an exploratory analysis of enhancers. According to the dbSUPER dabase only *CIPC* could work as a superenhancer in the brain middle hippocampus (SE_06106; chr14: 77562660-77607123, size: 44463 pb) (Supplementary Figure 3).

## CONCLUSION

As far as we know, this is the first study that has employed the UTMOST framework combined with the summary statistics of the largest ASD meta-analysis. The main aim was to identify ASD tissue-specific genes in brain and/or other tissues. It should be noted at the outset that gene-level associations identified by UTMOST do not imply causality. However, looking at the regional plots for each loci associated by UTMOST, the results shown at brain and gastrointestinal level seems pretty consistent. Thus, UTMOST has served to identify new ASD associated genes in brain tissue as *PTPRE* and *CIPC*. Furthermore, UTMOST has been useful to confirm and obtain information on the tissue in which other known ASD risk genes such as *NKX2-2* and P*INX1* may have functional relevance. Altogether, brain associated genes seems to point to three brain areas: cortex, hippocampus and nucleo accumbens. Previous studies demonstrated the ASD-associated gene expression in brain cortex^11^. In addition, hippocampus underlie some of the featured social memory and cognitive behaviors both crucial aspects in ASD ^12^. The nucleus accumbens is a key brain area also related with the social reward response in ASD^13^. It should be also noted some limitations of our study. UTMOST uses GTeX v6 data by default and it should be interesting to re-run the analysis once the tool will be updated. Thus, UTMOST could find novel associated genes when expression data from other relevant ASD brain tissues as the amygdala are included. Another limitation of our results is the small number of SNPs considered by UTMOST to create the statistic in some genes (*PTPRE* in brain tissues and *NKX2-2* in non-brain tissues analysis). However, we have been very conservative in establishing Bonferroni correction for all tissues in an analysis group. Thus, for example, we always chose the maximum number of genes tested in one of the brain tissues and applied this to calcule Bonferroni for the whole group of brain tissues as a whole.

In relation to the biological pathways in which UTMOST-associated genes are involved, our results open possible avenues for future genetic and functional studies. Thus, the functional role in ASD of *CIPC, PINX1, NKX2-2* and *PTPRE* has not yet been characterized in detail. Metascape analysis have provided an insight revealing their involvement in telomerase maintenance and transcription regulation. Thus, *PINX1* (*PIN2 (TERF1) Interacting Telomerase Inhibitor 1*) enhances TRF1 binding to telomeres and inhibits telomerase activity. It was proved that its silencing compromises telomere length maintenance in cancer cells ^14^. In addition, it was recently found that children with ASD and sensory symptoms have shorter telomeres, compared to those children exhibiting a typical development^15^. *CIPC* (*CLOCK interacting pacemaker*) is a mammalian circadian clock protein. Recently, circadian rithms were pointed out as involved in brain development and they could underly ASD etiology due to its implication in behavioral processes. Circadian rythms are regulated through several transcription factors in different cellular types, fact that might be related with the GO terms associated with transcription regulation in this study^16^. Moreover, the result of *CIPC* as a predicted superenhancer highlights its possible functional repercussion.

UTMOST found association of several genes in non-brain tissues. However, we found really interesting to study those genes related to gastrointestinal tissues (*NKX2-2, MANBA, ERI1* and *MITF*). There is a well-known and established comorbidity among gastrointestinal symptoms and ASD^17^. These clinical associations suggest the implication of gastrointestinal populations of neuronal cells. A mutation in *NLGN3* (R451C) has been recently identified in two ASD brothers with GI symptoms. Mice models have demonstrated that R451C alter the number of neuronal cells in the small intestine and impact fecal microbes^18^. These evidences suggest that the role of *NKX2-2, MANBA, ERI1* and *MITF* in gastrointestinal tissue should be further studied.

*BLK* and *NKX2-2* are both associated in the cross-tissue analysis. The importance of this approach is that is able to show the association of two previously known ASD risk genes but not restricted to brain tissue. These results lead us to a difficult question whether autism can be a multisystemic disorder, something that has been recently pointed out by some authors^19^.

In conclusion, UTMOST, a novel single and cross-tissue TWAS, has revealed new ASD associated genes. These genes have been characterized at the pathway and gene network level using bioinformatic approaches. However, future tissue-specific functional studies will be key to properly determine their role in ASD etiology.

## Supporting information

Supplementary .csv

Supplementary .csv

Supplementary .csv

Supplementary .csv

Supplementary .csv

Supplementary .csv

Supplementary .csv

Supplementary .csv

Supplementary .csv

Supplementary .csv

Supplementary .csv

Supplementary .csv

Supplementary .csv

Supplementary .csv

Supplementary .csv

Supplementary .csv

Supplementary .csv

Supplementary .csv

Supplementary .csv

Supplementary .csv

Supplementary .csv

Supplementary .csv

Supplementary .csv

Supplementary .csv

Supplementary .csv

Supplementary .csv

Supplementary .csv

Supplementary .csv

Supplementary .csv

Supplementary .csv

Supplementary .csv

Supplementary .csv

Supplementary .csv

Supplementary .csv

Supplementary .csv

Supplementary .csv

Supplementary .csv

Supplementary .csv

Supplementary .csv

Supplementary .csv

Supplementary .csv

Supplementary .csv

Supplementary .csv

Supplementary .csv

## ACKNOWLEDGEMENTS

We thank the Psychiatric Genomics Consortium Autism Spectrum Disorder Working Group for making the ASD genome-wide association study results publicly available. The authors report no biomedical financial interests or potential conflicts of interest.

## WEB LINKS AND URLS

The supplementary material for this article can be found online at:

---

**Supplementary Figure 1a.**
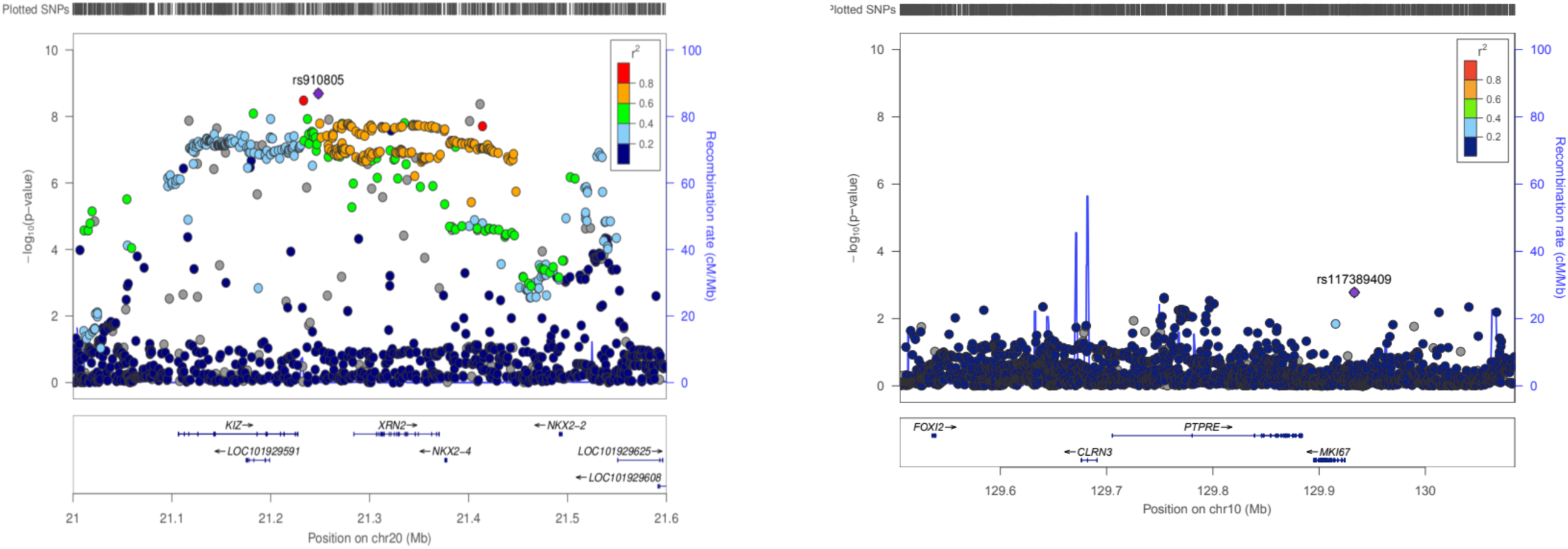
Regional plots using ASD metaanalysis summary statistics for the UTMOST associated genes in brain tissues (*NKX2-2, PTPRE*).

**Supplementary Figure 1b.**
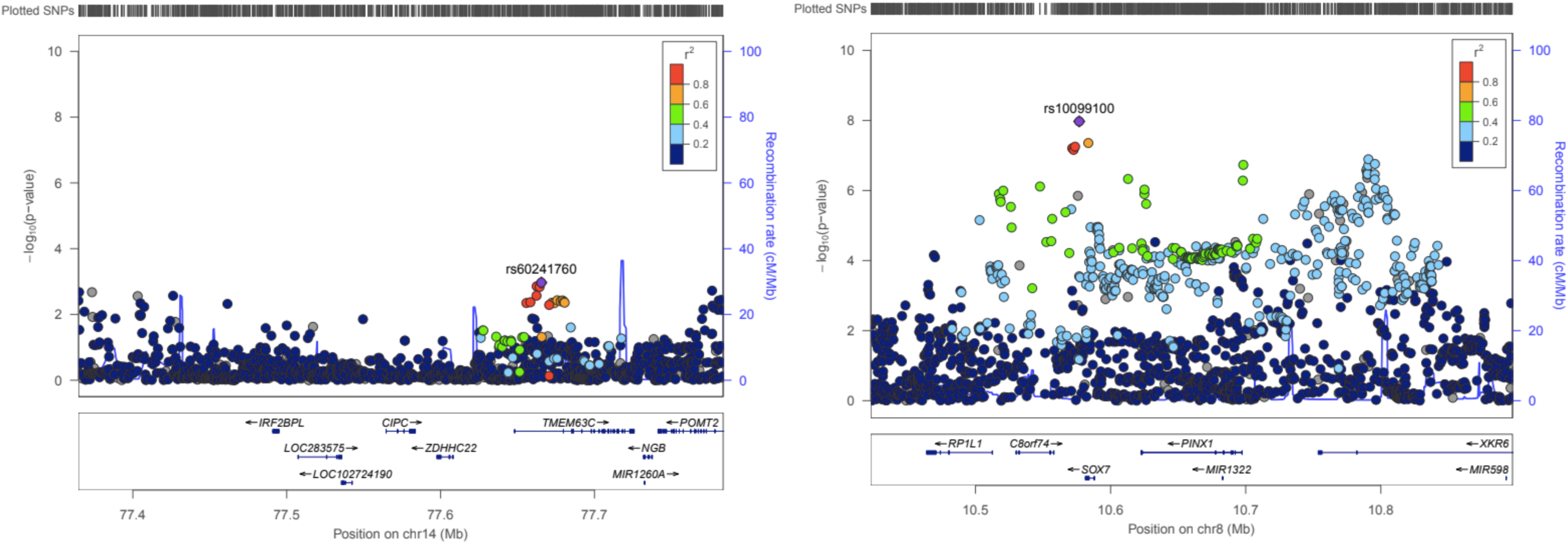
Regional plots using ASD metaanalysis summary statistics for the UTMOST associated genes in brain tissues (*CIPC, PINX1*).

**Supplementary Figure 2a.**
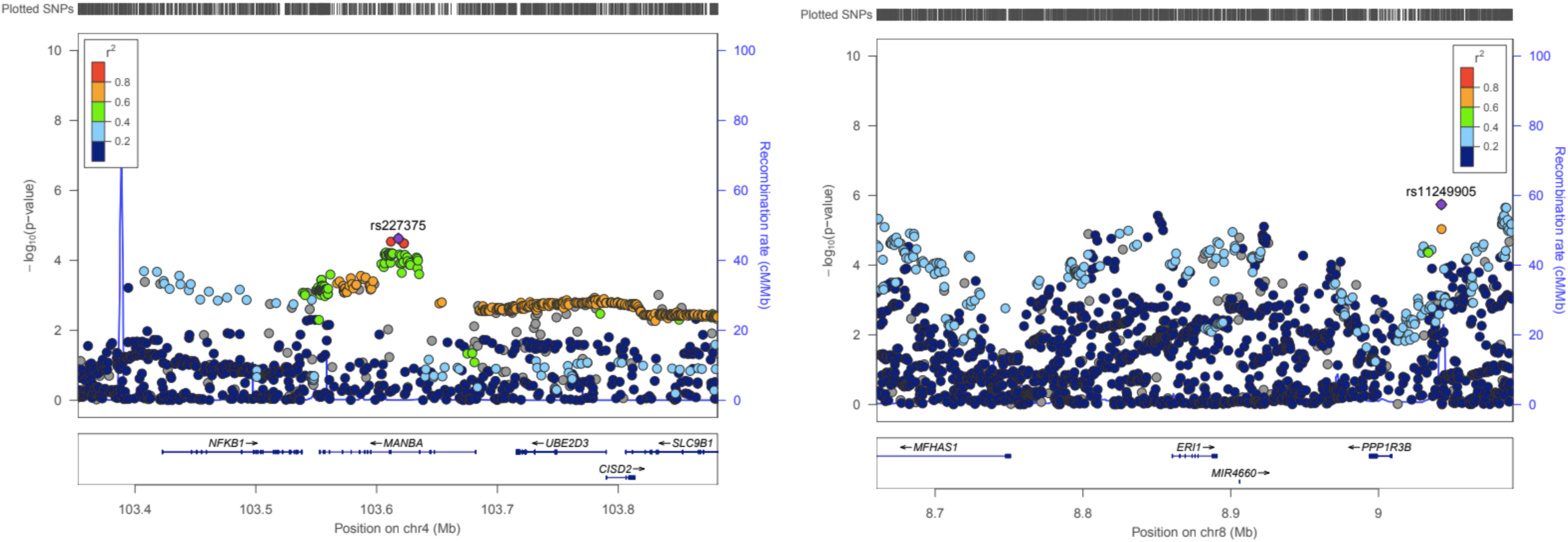
Regional plots using ASD metaanalysis summary statistics for the UTMOST associated genes in gastrointestinal tissues (*MANBA, ERI1*). *NKX2-2* has shown association in this analysis but the regional plot is included in the previous Supplementary Figure 1.

**Supplementary Figure 2b.**
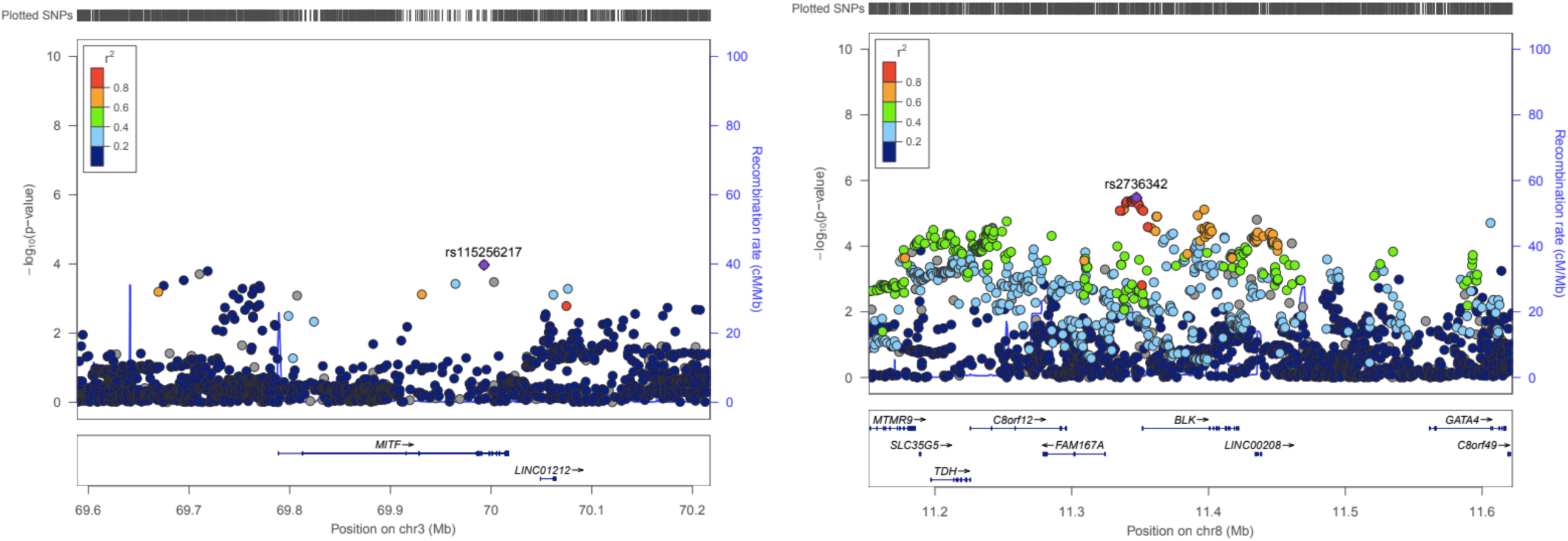
Regional plots using ASD metaanalysis summary statistics for the UTMOST associated gene in gastrointestinal tissues *MITF* and cross-tissue analysis *BLK*. *NKX2-2* is also associated in these analysis but the regional plot is included in the Supplementary Figure 1.

**Supplementary Figure 3.**
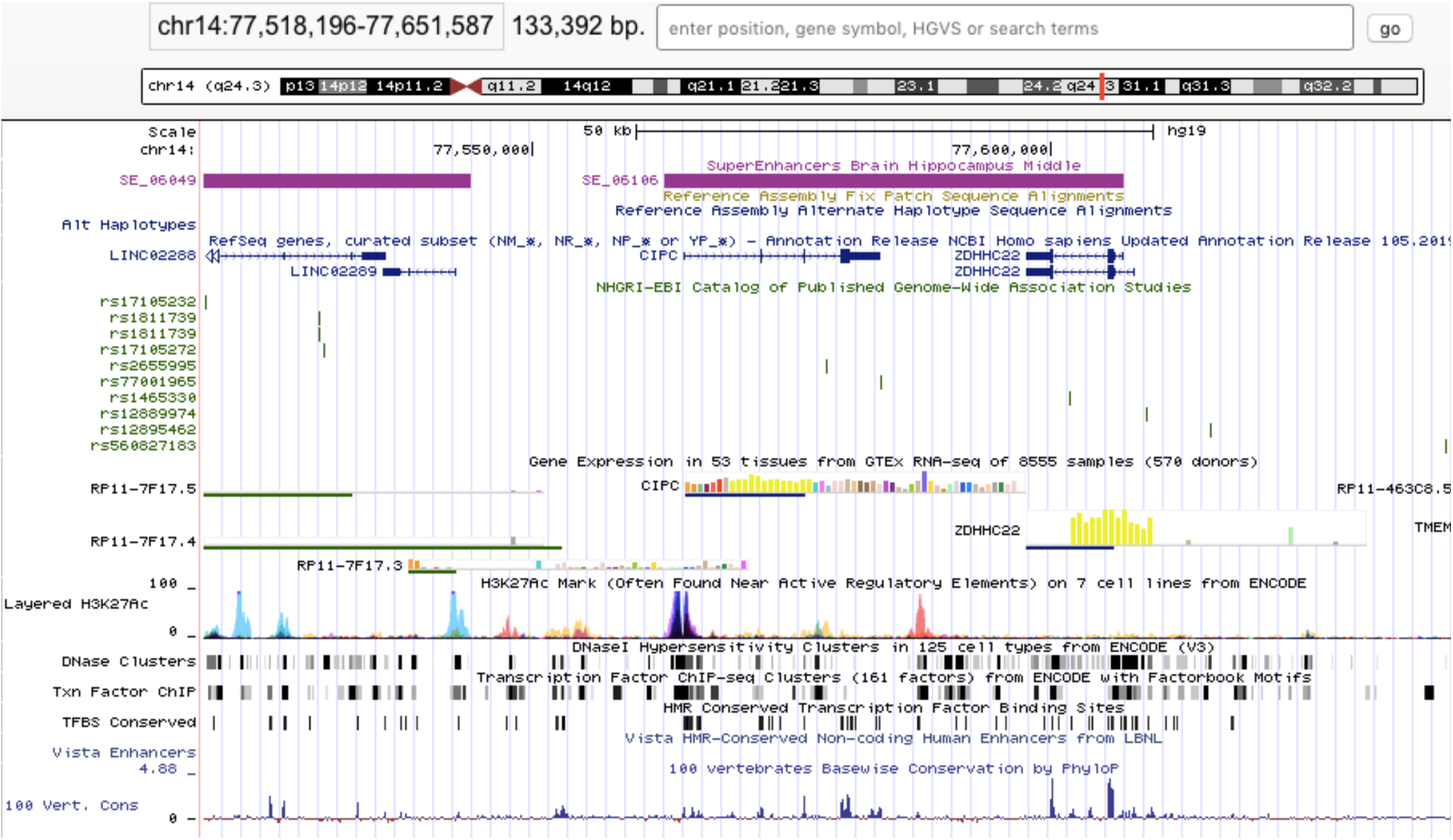
UCSC screenshot showing the location of SE_06106 superenhancer on CIPC gene (hippocampus).

## REFERENCES

1. American Psychiatric Association. Diagnostic and Statistical Manual of Mental Disorders (Arlington, VA: American Psychiatric Publishing). (2013).

2. Sanders, S. J. et al. Insights into Autism Spectrum Disorder Genomic Architecture and Biology from 71 Risk Loci. Neuron 87, 1215–1233 (2015).

3. Grove, J. et al. Identification of common genetic risk variants for autism spectrum disorder. Nat. Genet. 51, 431–444 (2019).

4. Alonso-Gonzalez, A., Calaza, M., Rodriguez-Fontenla, C. & Carracedo, A. Novel Gene-Based Analysis of ASD GWAS: Insight Into the Biological Role of Associated Genes. Front. Genet. 10, (2019).

5. Barbeira, A. N. et al. Exploring the phenotypic consequences of tissue specific gene expression variation inferred from GWAS summary statistics. Nat. Commun. 9, 1825 (2018).

6. Gusev, A. et al. Integrative approaches for large-scale transcriptome-wide association studies. Nat. Genet. 48, 245–252 (2016).

7. Hu, Y. et al. A statistical framework for cross-tissue transcriptome-wide association analysis. Nat. Genet. 51, 568–576 (2019).

8. Watanabe, K., Taskesen, E., van Bochoven, A. & Posthuma, D. Functional mapping and annotation of genetic associations with FUMA. Nat. Commun. 8, 1826 (2017).

9. Warde-Farley, D. et al. The GeneMANIA prediction server: biological network integration for gene prioritization and predicting gene function. Nucleic Acids Res. 38, W214–220 (2010).

10. Zhou, Y. et al. Metascape provides a biologist-oriented resource for the analysis of systems-level datasets. Nat. Commun. 10, 1523 (2019).

11. Voineagu, I. et al. Transcriptomic analysis of autistic brain reveals convergent molecular pathology. Nature 474, 380–384 (2011).

12. Hitti, F. L. & Siegelbaum, S. A. The hippocampal CA2 region is essential for social memory. Nature 508, 88–92 (2014).

13. Park, H. R. et al. A Short Review on the Current Understanding of Autism Spectrum Disorders. Exp. Neurobiol. 25, 1–13 (2016).

14. Zhang, B. et al. Silencing PinX1 compromises telomere length maintenance as well as tumorigenicity in telomerase-positive human cancer cells. Cancer Res. 69, 75–83 (2009).

15. Lewis, C. R. et al. Telomere Length and Autism Spectrum Disorder Within the Family: Relationships With Cognition and Sensory Symptoms. Autism Res. n/a,.

16. Geoffray, M.-M., Nicolas, A., Speranza, M. & Georgieff, N. Are circadian rhythms new pathways to understand Autism Spectrum Disorder? J. Physiol. Paris 110, 434–438 (2016).

17. Buie, T. et al. Evaluation, diagnosis, and treatment of gastrointestinal disorders in individuals with ASDs: a consensus report. Pediatrics 125 Suppl 1, S1–18 (2010).

18. Hosie, S. et al. Gastrointestinal dysfunction in patients and mice expressing the autism-associated R451C mutation in neuroligin-3. Autism Res. 12, 1043–1056 (2019).

19. Thom, R. P. et al. Beyond the brain: A multi-system inflammatory subtype of autism spectrum disorder. Psychopharmacology (Berl.) 236, 3045–3061 (2019).

